# Distinct activation thresholds of unmyelinated C-fiber afferents by dorsal root ganglion and peripheral nerve stimulation

**DOI:** 10.64898/2026.02.02.703367

**Authors:** Longtu Chen, Jia Liu, Shaopeng Zhang, Sajjad Rigi Ladez, Bin Feng

## Abstract

**Objectives:** Sensitization of C-fiber nociceptors plays a critical role in spontaneous and ongoing pain in patients with chronic pain. We recently demonstrated that C-fiber afferents can be reversibly blocked through activity-dependent conduction slowing, suggesting that selective activation of C-fiber afferents may represent a novel strategy for pain relief. We hypothesized that electrical peripheral nerve stimulation (ePNS) and dorsal root ganglion (DRG) stimulation exhibit distinct activation thresholds for unmyelinated C-fiber afferents.

**Materials and Methods:** We characterized the activation thresholds of A- and C-fiber afferents during ePNS and DRG stimulation using single-fiber electrophysiological recordings from split nerve filaments and optical GCaMP6f imaging of intact DRGs. We further quantified the distribution of the sodium channel subtype NaV1.6 in afferent axons and somata and evaluated its contribution to neuronal excitability using NEURON-based computational modeling.

**Results:** Single-fiber recordings showed that activation of C-fiber axons required approximately tenfold higher stimulus amplitudes than Aα/β-fiber axons during ePNS. In contrast, DRG stimulation within a narrow amplitude range robustly activated both small- and large-diameter DRG neurons, with putative C-fiber afferents comprising 57% of the activated population, indicating markedly reduced differential activation thresholds compared with ePNS. Analysis of published single-cell RNA-sequencing datasets revealed high NaV1.6 expression in TRPV1-positive C-fiber nociceptors. Immunohistochemical staining demonstrated prominent clustering of NaV1.6 in the stem axons of most DRG neurons, including small-diameter C-fiber afferents, whereas NaV1.6 was absent from C-fiber axons in the sciatic nerve. NEURON simulations further showed that NaV1.6 clustering at the stem axon is a key determinant of activation thresholds during DRG stimulation.

**Conclusions:** These findings identify a structural and molecular mechanism underlying the efficient activation of C-fiber afferents by DRG stimulation and provide mechanistic insight into the superior therapeutic efficacy of DRG stimulation for the treatment of C-fiber–mediated chronic pain.

## Introduction

Electrical stimulation of peripheral sensory afferents has been extensively applied in the clinic for managing a variety of disorders, including electrical peripheral nerve stimulation (ePNS) targeting sensory axons in nerve trunks and dorsal root ganglion (DRG) stimulation targeting afferent somata. ePNS with electrodes placed outside the epineurium of peripheral nerve trunks has been widely used for treating chronic pain ^1^, fecal/urinary incontinence ^2, 3^, and to a lesser extent, obesity ^4^. Invasive placement of leads inside the nerve trunk, e.g., with transverse intrafascicular multichannel electrode (TIME), longitudinal intrafascicular electrode (LIFE), and Utah slanted electrode array (USEA), has limited clinical applications largely due to concerns about device longevity ^5^. The treatment of chronic pain by ePNS is generally attributed to the activation of myelinated A-fiber afferents to inhibit nociceptive transmission in the spinal cord, according to the “Gate Control Theory ^6^”. Activation of large A-fiber axons at low frequencies (usually less than 100 Hz) evokes paresthesia, a non-painful tingling sensation that blocks nociceptive signaling from the same area ^7^. This mechanism of action contrasts with chemical nerve blocks, e.g., with lidocaine, which suppress pain by blocking conduction in both myelinated A-fiber and unmyelinated C-fiber axons ^8^. Epineural ePNS reportedly activates large, myelinated A-fiber axons at a much lower stimulus threshold than small unmyelinated C-fiber axons ^9^. Accordingly, the neuromodulatory target of ePNS is predominantly myelinated A-fiber axons. Since blocking neural transmission of large-diameter A-fiber axons requires kilohertz electrical stimulation ^10, 11^, ePNS that utilizes nerve block for treating obesity delivers kilohertz stimulation to the vagal nerve ^4^.

The sensory afferent somata in the dorsal root ganglia (DRG) have also been targeted by electrical neuromodulation. DRG stimulation was approved by the FDA for treating complex regional pain syndrome in the lower limb in 2016 ^12^. Compared to ePNS, DRG stimulation requires more invasive placement of electrode leads in contact with the epidural surface of the ganglia ^13^, and is less widely used in the clinic. Recent clinical observations indicate that DRG stimulation can relieve pain in patients without causing paresthesia ^14^. Compared with patients experiencing paresthesia, paresthesia-free patients reported comparable and even superior therapeutic outcomes from DRG stimulation ^15^. We and others have demonstrated that C-fiber afferents can be reversibly blocked by sub-kilohertz DRG stimulation (usually <100 Hz), which provides a mechanistic interpretation of pain suppression in the absence of paresthesia ^16, 17^. Recently, we overcame the limitation of activating C-fiber axons with ePNS by implementing a liquid suction electrode to increase the seal resistance between the electrode and surrounding solution ^9^. With this approach, we reported that sub-kilohertz ePNS reversibly blocked action potential transmission in C-fiber axons, similar to DRG stimulation ^9^.

Electrophysiological studies suggest differential activation of A-fiber and C-fiber afferent populations by stimulating the DRG and peripheral nerve trunk. However, the exact thresholds for activating different types of afferents with ePNS and DRG stimulation have not been systematically determined and reported. No prior studies have provided speculations or hypotheses on the underlying mechanisms that account for this differential activation of afferent populations by ePNS and DRG stimulation. By analyzing published single-cell RNA sequencing databases ^18, 19^, we found that over 43.78% of DRG neurons with markers for unmyelinated C-fibers express NaV1.6, a voltage-gated sodium channel subtype known for its low activation thresholds, rapid activation and inactivation, and the presence of persistent and resurgent currents ^20^. NaV1.6 is widely expressed in myelinated neurons that fire at high frequencies and/or repetitively, for example, Purkinje neurons in the cerebellum ^21^ and muscle spindle afferents ^22^. Regarding unmyelinated C-fiber afferents, we previously reported that NaV1.6 plays necessary roles in the tonic spiking of stretch-sensitive visceral afferent endings innervating the distal large intestine ^23^, which are predominantly unmyelinated C-fibers ^24^. The experimental evidence collectively suggests that the differential distribution of NaV1.6 in afferent axons and somata likely contributes to their different excitability to extracellular electrical stimulation.

In this study, we aim to systematically determine the differential activation of both A-fiber and C-fiber afferents by ePNS versus DRG stimulation. We determined the activation threshold by conducting single fiber recordings from split peripheral nerve filaments and optical GCaMP6f recordings from individual DRG neurons. We also investigated the clustering of NaV1.6 by immunohistological staining at afferent axons, DRG somata, and stem axons. Additionally, we conducted computational NEURON simulations to systematically assess the impact of NaV1.6 clustering in the stem axon of unmyelinated C-fiber afferents on the activation threshold.

## Materials and Methods

All experiments were reviewed and approved by the University of Connecticut Institutional Animal Care and Use Committee. Mice were housed in pathogen-free facilities accredited by the American Association for Accreditation of Laboratory Animal Care and assured by the Public Health Service, following guidelines set forth in the Eighth Edition of the Guide for the Care and Use of Laboratory Animals. Housing consisted of individually ventilated polycarbonate cages (Animal Care System M.I.C.E.) with a maximum occupancy of 5 mice per cage. Environmental enrichment included nestlets and huts, with Envigo T7990 B.G. Irradiated Teklad Sani-Chips as bedding material. Mice were fed ad lib with either 2918 Irradiated Teklad Global 18% Rodent Diet or 7904 Irradiated S2335 Mouse Breeder Diet supplied by Envigo and supplied with reverse osmosis water chlorinated to 2 ppm using a water bottle. The animal housing facility maintained a 12:12 light-dark cycle, with ambient temperature regulated between 70-77°F (set point 73.5°F) and relative humidity between 35-65% (set point 50%). Animal care services staff performed daily health checks, and cages were changed biweekly.

### Ex vivo single-fiber recordings from split sciatic nerve axons

From adult C57BL/6 mice of both sexes (aged 10-16 weeks, weighing 25-35 g), we harvested the sciatic nerve and its most distal branch, tibial nerve following surgical procedures previously reported ^25, 26^. Briefly, mice were anesthetized with 2% isoflurane inhalation and euthanized by transcardiac perfusion with oxygenated Krebs solution (in mM: 117.9 NaCl, 4.7 KCl, 25 NaHCO_3_, 1.3 NaH_2_PO_4_, 1.2 MgSO_4_, 2.5 CaCl_2_, and 11.1 D-glucose at room temperature) from the left ventricle to the right atrium through the circulatory system. The perfused carcass was then immediately transferred to a dissection chamber circulated with oxygenated ice-cold Krebs solution for nerve harvesting. The sciatic nerve, approximately 35-40 mm in length, was meticulously harvested from its proximal projection at the L4 spinal cord to its distal projection at the tibial nerve in the heel.

As shown in **Fig. 1A**, harvested nerves were then transferred to a custom-built two-compartment chamber consisting of a tissue compartment and a recording compartment ^25^. The proximal end of the targeted nerve was pinned down in the tissue compartment circulated with oxygenated Krebs solution at 28 to 30°C. The ∼5 mm distal end of the targeted nerve was gently pulled over and laid onto a mirror in the recording compartment filled with mineral oil (Fisher Scientific, East Greenwich, RI) to enhance the signal-to-noise ratio of the recording. Then, the distal end of the nerve (recording chamber side) was split into fine filaments (∼10 µm thickness) to achieve optimal single-fiber recordings from individual afferent/efferent axons ^25, 26^. A customized 5-channel electrode array was utilized to interface with split nerve filaments ^25^. Single-unit action potentials (APs) from all five electrodes were recorded simultaneously, digitized at 20 kHz and band-pass filtered (200–3000 Hz) using an Intan RHD USB interface board. The multichannel recording signals were also monitored by a data acquisition system (1401plus, CED, Cambridge, UK) and stored onto a PC using Spike2 software (v7.1, CED, Cambridge, UK).

**Figure 1.**
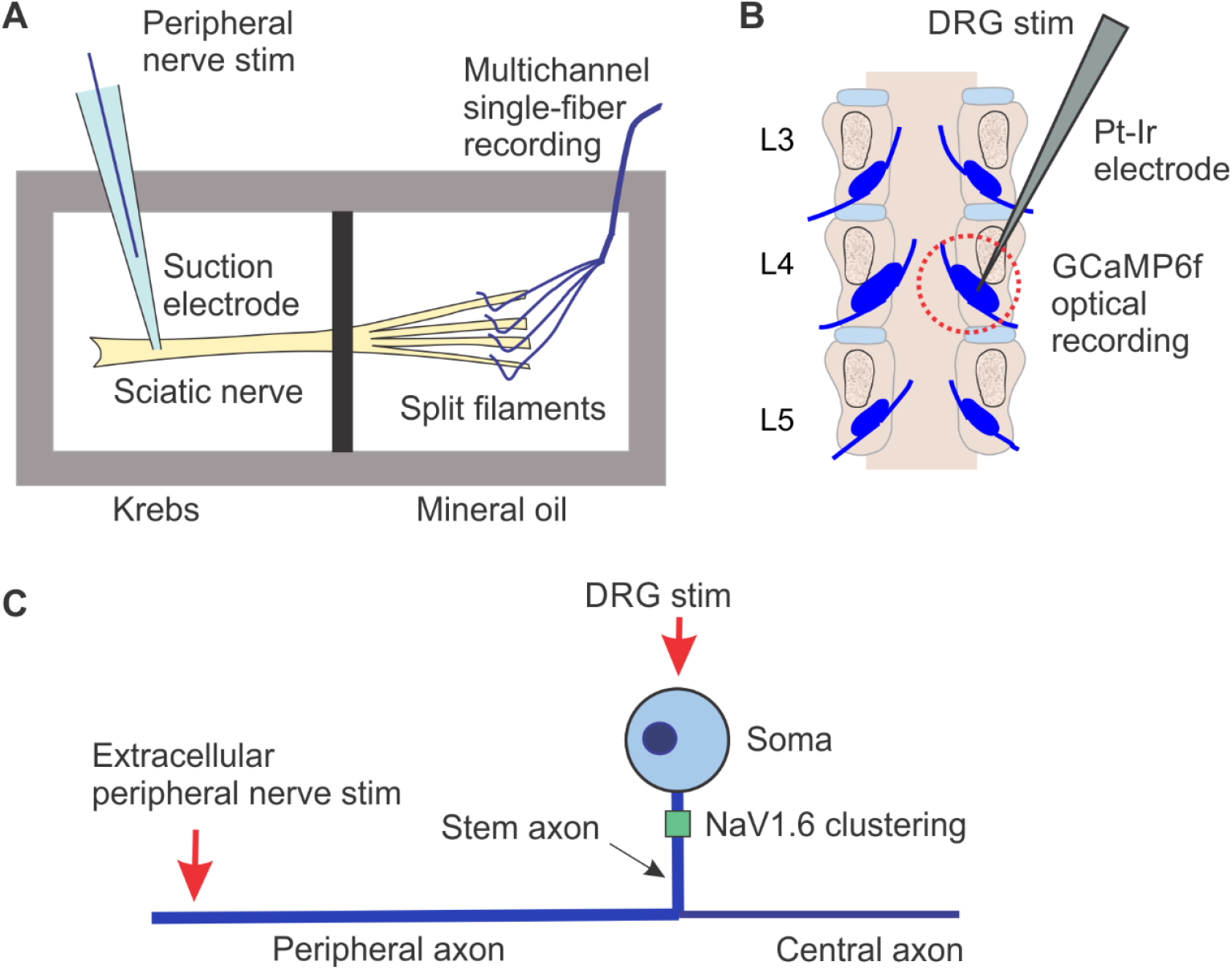
Schematic diagrams illustrating (A) single-fiber recordings from split sciatic nerve filaments used to determine ePNS activation thresholds; (B) GCaMP6f imaging of DRG somata used to assess DRG stimulation thresholds; and (C) the corresponding NEURON simulation model of an unmyelinated C-fiber afferent.

As reported previously ^9^, action potentials (APs) were evoked at the proximal end of the sciatic nerve using a custom-built liquid suction electrode, created by pulling quartz glass capillaries in a micropipette puller (P-97, Sutter Instrument, Novato, CA) to form a tip of ∼Φ400 µm, approximately 30% smaller than the nerve diameter. The suction electrode, filled with Krebs solution, formed a loose seal with the epineurium surface via gentle negative pressure (-60 to -30 mmHg). Care was taken to clean surrounding connective tissue to aid seal formation, which is crucial for evoking unmyelinated C-fiber axons ^9^. APs were evoked at 0.5 Hz using monophasic constant current stimulation of 0.2 msec pulse width (cathodic, A-M Systems 4100, Carlsborg, WA). The activation threshold was determined at the pulse amplitude to evoke 4 to 6 APs per 10 stimuli.

### Ex vivo GCaMP6f recordings from whole DRG

To optically record the afferent neuron’s activation threshold by DRG stimulation, we used transgenic mice expressing the fluorescent calcium indicator GCaMP6f in glutamatergic neurons as reported previously ^16, 27, 28^. Ai95 mice (C57BL/6 background) possessing homozygous GCaMP6f (strain# 28865, Jackson Laboratory, CT) were crossbred with mice carrying homozygous VGLUT2-Cre (strain# 28863, Jackson Laboratory, CT)^16^. We used transgenic mice of both sexes (8–16 weeks, 25–35 g) with expression of GCaMP6f genes for optical recordings. Following the same euthanasia and transcardiac perfusion procedures described above, a dorsal pediculectomy was performed to expose the spinal cord and DRG. The L3, L4, and L5 DRG that receive spinal nerve projections were harvested and placed in a tissue chamber perfused with oxygenated Krebs solution at 30°C, as illustrated in **Fig. 1B**. Optical GCaMP6f recordings were conducted using an upright fluorescent microscope (BX51WI, Olympus, Waltham, MA) equipped with a water immersion 20x objective (UMPLFLN 10XW, 0.3 NA). This setup allowed visualization of the entire DRG with sufficient resolution to detect calcium transients in individual DRG somata. The DRG was illuminated using a halogen epi-illumination light source. GCaMP6f transients were recorded using a high-speed, ultra-low noise sCMOS camera (Xyla-4.2P, 82% quantum efficiency, Andor Technology, South Windsor, CT). Post-hoc analysis of DRG dimensions was performed on captured images using ImageJ (NIH, Bethesda, MD).

To electrically stimulate the DRG, we used a blunt-tipped needle electrode (FHC, platinum-iridium, tip size ∼Φ5 μm) placed in contact with but not penetrating the dura mater of the DRG to deliver constant current stimulation (701C stimulating module, Aurora Scientific Inc., Canada). A monopolar stimulation was configured with the counter electrode placed in the Krebs bath solution. Charge-balanced biphasic stimuli (constant current, cathodic first) were delivered at 0.5 Hz with a 0.2 msec pulse phase width and 0.05 msec interphase interval (IPI). The activation threshold was determined at the pulse amplitude to evoke 4 to 6 GCaMP6f fluorescent signals per 10 stimuli. To avoid prolonged characterization that could result in fluorescence photobleaching, activation thresholds were evaluated over a range of 0 to 0.5 mA with 0.05 mA intervals.

### Fluorescent labeling of afferents

We utilized Cre-dependent adeno-associated viruses (AAVs) for sparse labeling of afferent somata and axons in VGLUT2-Cre mice (strain#. 28863, The Jackson Laboratory, CT) using AAV9-floxed-ChR-EYFP (#20298-AAV9, Addgene). We implemented the EYFP fused with Channelrhodopsin2 (ChR-EYFP) as the fluorescent reporter for our anatomic tracing. This fluorescent reporter serves as a membrane-bound marker and has been demonstrated to transport over 10 mm from the somata to peripheral nerve endings in previous optogenetic studies on the dorsal root ganglia ^29-31^. Mice were anesthetized by inhalation of 1.5% isoflurane, and a small (∼1 cm) midline incision of the skin was made over the T12-L1 vertebra to expose the dorsal tissue. Muscles and tendons were severed to expose the dura mater between the T13 and L1 vertebrae. A 33-gauge needle (model 7643-01, Hamilton Company, Arlington, MA) was inserted into the intrathecal space, and 5 μL AAV9-floxed-ChR-EYFP (titer >1×10^13^ vg/mL) was slowly injected. After injection, the skin was sutured with non-absorbable polypropylene suture (8698G, Ethicon). Meloxicam (2 mg/kg, Boehringer Ingelheim Vetmedica, Duluth, GA) was given three times in 72 hours for postoperative analgesia. After injection, mice were allowed 6 weeks for sufficient AAV transfection.

In some experiments designed to assess neural structures near DRG somata, afferents were labeled by crossbreeding VGLUT2-Cre mice with tdTomato LoxP reporter mice (strain# 007905, The Jackson Laboratory, CT), an approach that fluorescently labeled the intracellular space of afferent somata and stem axons in the DRG ^31^.

### Immunohistological staining of DRG, afferent axons in spinal nerves, and afferent endings

Mice were deeply anesthetized and euthanized by transcardiac perfusion from the left ventricle with ice-cold oxygenated Krebs solution as described above. The DRGs (T12-S1), sciatic nerve, and bladder were harvested from the mouse and immediately placed in ice-cold oxygenated Krebs solution. The bladder was cut open, slightly stretched axially and circumferentially, and pinned flat in the Sylgard-lined Petri dish. All the harvested tissue was then fixed with 10% buffered formalin at 4 °C overnight. The fixed tissue was washed with phosphate-buffered saline (PBS) three times (10 min each) to remove the fixative, cryoprotected in 20% sucrose in PBS, embedded in OCT compound (Sakura Finetek, Tokyo, Japan), frozen, and sectioned at 40 µm. Tissue sections were incubated with rabbit antibodies against NaV1.6 (1:1000, Alomone) and co-stained with rat antibodies against GFP (1:1000, MBL International Corp) for 24 h at 4 °C, and then incubated with a mixture of secondary antibodies Alexa Fluor® 594-conjugated anti-rabbit IgG (1:200, Abcam) and Alexa Fluor® 488-conjugated anti-rat IgG (1:200, Abcam) for 2 h at room temperature. Afterwards, the stained sections were mounted in reflective index matching solution for subsequent microscopic imaging. Fluorescent images were analyzed using ImageJ (NIH, Bethesda, MD) and data were presented with box-and-whisker plots with overlaid scatter plots.

### Computational modeling study of afferent activation by ePNS and DRG stimulation

Using the NEURON simulation environment ^32^, we developed a biophysically detailed model of a mouse unmyelinated C-fiber afferent to systematically examine the impact of NaV1.6 clustering in the stem axon on activation thresholds during extracellular ePNS and DRG stimulation. As illustrated in **Fig. 1C**, the model comprised a 23-µm soma connected to a 100-µm stem axon, which bifurcated into central (2500 µm) and peripheral (5000 µm) projections. A 4-µm segment of the stem axon, located 6.4 µm from the soma, contained a concentrated distribution of NaV1.6 channels, consistent with the immunohistological staining pattern observed in this study. The diameters of the stem and peripheral axons were set to 1.2 µm, and the central axon to 1.0 µm.

Biophysical parameters were adopted from our prior work ^9, 23^. Briefly, the intracellular resistivity (Ra) was set to 123 Ω·cm and the membrane capacitance to 1 µF/cm^2^. Active membrane mechanisms included voltage-gated sodium channels (NaV1.6, NaV1.7, NaV1.8, NaV1.9), potassium channels (fast and intermediate inactivating [K_A_], slow inactivating [K_D_], and delayed rectifier [K_S_]), the Na^+^/K^+^-ATPase (NKA), and a passive leak conductance. Channel kinetics were implemented using Hodgkin–Huxley-type or Markov-state models as summarized in **Table 1**.

**Table 1.** Transmembrane ionic conductance in the model.

| Conductance | Model type | Maximum conductance ($\text{S}/\text{cm}^2$) | | |
| --- | --- | --- | --- | --- |
|  |  | Soma | Axons | Axons with NaV1.6 |
| NaV1.6 | Markov | 0 | 0 | 2 |
| NaV1.7 | Markov | 0.008 | 0.2 | 0.2 |
| NaV1.8 | HH-type | 0.02 | 0.5 | 0.5 |
| NaV1.9 | HH-type | 0.0001 | 0.0002 | 0.0002 |
| $K_A$ | HH-type | 0.09 | 0.045 | 0.045 |
| $K_D$ | HH-type | 0.08 | 0.04 | 0.04 |
| $K_S$ | HH-type | 0.038 | 0.021 | 0.021 |
| NKA |  | 0.0025 | 0.0025 | 0.0025 |
| Leak |  | 0.001 | 0.001 | 0.001 |

Extracellular stimulation was implemented in NEURON using the built-in extracellular mechanism with a point-source current injection. As shown in **Fig. 1C**, the point source was positioned 5 µm above the soma to model DRG stimulation, and 5 µm lateral to a site located 100 µm from the distal end of the peripheral axon to simulate ePNS. To match experimental conditions, monophasic ePNS pulses (0.2 ms) were delivered to the peripheral axon at 0.5 Hz, and biphasic DRG stimuli (0.2 ms, cathodic-first, 0.05 ms inter-phase interval) were applied to the soma at 0.5 Hz. Action-potential thresholds were assessed both at the spike-initiation site nearest the point source and at the distal ends of the peripheral and central axons to confirm successful propagation. To systematically evaluate how NaV1.6 clustering affects afferent excitability, activation thresholds were quantified across a range of maximum NaV1.6 conductances within the clustered segment of the stem axon. All simulations were performed at 30 °C, consistent with the conditions used in the ex vivo single-fiber recordings

### Data Analysis and Statistics

Action potentials recorded by single-fiber recordings were processed off-line using customized MATLAB program. The detection thresholds for individual action potentials were set as four times the root mean square (RMS) amplitude of background noise recorded 10 msec before the stimulation. Conduction delays were measured as the time between the onset of stimulus artifacts and the onset of recorded action potentials. The conduction velocity (CV) was computed from the conduction delay and the distance between the stimulation and the recording sites. From each harvested sciatic nerve, action potentials from 4 to 6 axons were recorded, including at least one A-fiber axon with CV greater than 1 m/s. To facilitate between-sample comparison, activation thresholds recorded within each sciatic nerve were normalized by the lowest threshold. For GCaMP6f recordings, DRG neurons were grouped by soma size into small-(Φ < 27 µm) and large-diameter groups (Φ ≥ 27µm), which generally correlate with C- and A-type DRG neurons ^33, 34^. DRG diameters were quantified by measuring the diameter of the smallest enclosing circle using ImageJ ^35^. DRG neurons were considered activated when exhibiting an increase of GCaMP6f fluorescent signals greater than 10% of the baseline fluorescent signal (*ΔF/F* > 10%). Data are presented as means ± standard error (S.E.). One-way ANOVA was performed as appropriate using SigmaStat v4.0 (Systat Software, San Jose, CA). Differences were considered significant when p < 0.05.

## Results

### Significant presence of NaV1.6 in mouse DRG neurons

Analysis of single-cell RNA sequencing (scRNAseq) data from published databases revealed distinct expression patterns of NaV1.6 across different dorsal root ganglion (DRG) neuron populations as quantified by reads per million (RPM) (**Fig. 2**). There are four main identified neuronal clusters based on their expression of known markers (**Fig. 2A**) ^19^. The NF (neurofilament) cluster, expressing Nefh and Pvalb, is associated with myelinated DRG neurons. The PEP (peptidergic) cluster, expressing Tac1, Ntrk1, and Calca, is associated with peptidergic nociceptors. The NP (non-peptidergic) cluster, expressing Mrgprd and P2rx3, is associated with non-peptidergic nociceptors. Lastly, the TH (tyrosine hydroxylase) cluster, distinctly expressing Th, is associated with a subclass of unmyelinated neurons. The expression of NaV1.6 (Scn8a) was observed across all these clusters. Over 90.65% of DRG neurons with markers for myelinated A-fibers (primarily in the NF cluster) expressed Scn8a. Surprisingly, 43.78% of DRG neurons with markers for unmyelinated C-fibers (primarily in the PEP, NP, and TH clusters) also expressed Scn8a as indicated with Log(RPM) greater than 0 (RPM >1) in **Fig. 2A**.

**Figure 2.**
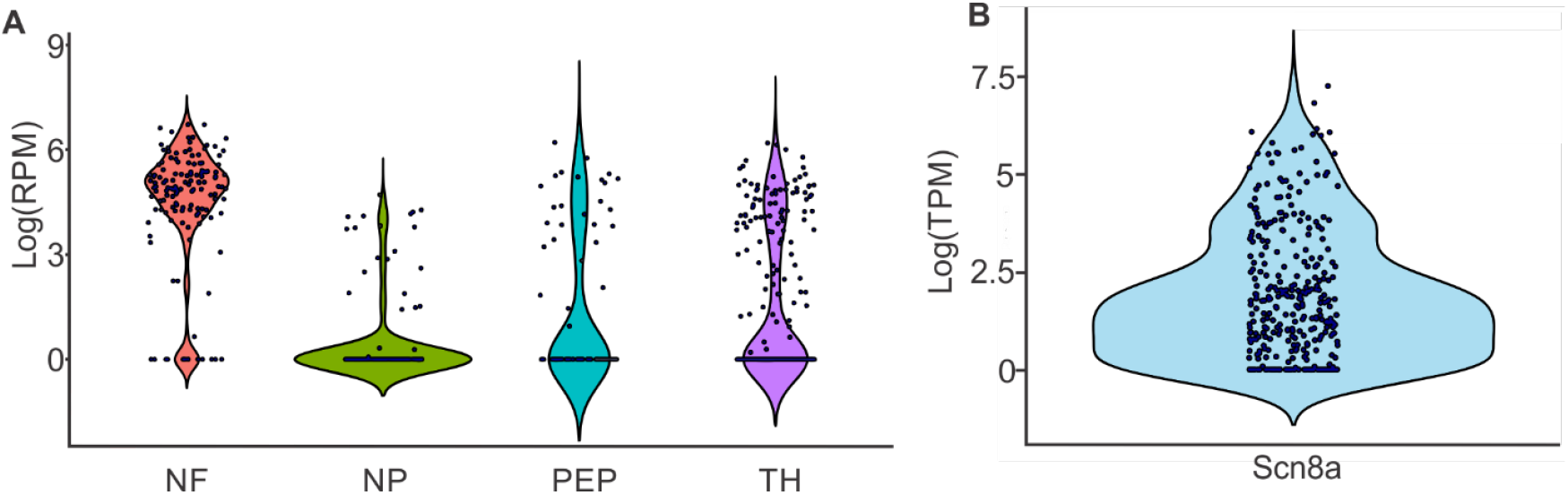
Analyses of two prior single-cell RNA-seq studies of mouse DRG neurons reveal widespread expression of NaV1.6 across both A- and C-fiber afferents ^18, 19^. (A) Violin plots showing NaV1.6 (Scn8a) mRNA expression levels across distinct DRG neuronal populations. NF, neurofilament; NP, non-peptidergic; PEP, peptidergic; TH, tyrosine hydroxylase. (B) Violin plot showing NaV1.6 mRNA expression in colonic DRG neurons. RPM, reads per million; TPM, transcripts per million.

Furthermore, analysis of another database, focusing specifically on colonic DRG neurons, showed that 81.75% of these neurons have substantial expression of NaV1.6 mRNA, as quantified by transcripts per million (TPM) with Log(TPM) greater than 0 (TPM >1) (**Fig. 2B**) ^18^. Our prior functional characterization study reported that most, if not all colonic DRG neurons are unmyelinated C-fibers with CV less than 1 m/s ^24^. This widespread expression of NaV1.6, the voltage-gated sodium channel subtype known for its low activation thresholds, rapid activation and inactivation, and the presence of persistent and resurgent currents, suggests its potential importance in regulating the excitability of both A-fiber and C-fiber DRG neurons.

### Differential afferent activation thresholds between ePNS and DRG stimulation

Displayed in **Fig. 3A** are typical single-fiber recordings of evoked APs from ePNS. At low-intensity stimulation, ePNS evokes exclusively fast-conducting A-fiber axons with CV > 1 m/s. With increased stimulus intensity, slow-conducting C-fiber axons (CV < 1 m/s) were also recruited. Significantly higher stimulus artifacts were evoked by high-intensity stimulation. In **Fig. 3B** are representative GCaMP6f recordings of evoked afferent neurons by DRG stimulation, showing that small-diameter neurons (putative C-fiber afferents) can be evoked at lower stimulus intensity (0.3 mA) than large-diameter neurons (0.45 mA). As summarized in **Fig. 3C**, activation thresholds by ePNS were determined from a total of 27 afferents with different CV, which were recorded from 4 different sciatic nerves. Thresholds of afferents recorded from the same nerve were normalized by the lowest threshold to facilitate between-sample comparison. Based on their CV, afferents were categorized as C-fibers (CV < 1m/s, N = 6), Aδ-fibers (1 m/s ≤ CV < 7 m/s, N = 13), and Aα/β-fibers (CV ≥ 7 m/s, N = 8). Displayed in **Fig. 3D** are activation thresholds by DRG stimulations, which were determined from 30 DRG neurons with different diameters (Φ < 27 μm, N = 17; Φ ≥ 27 μm, N = 13).

**Figure 3.**
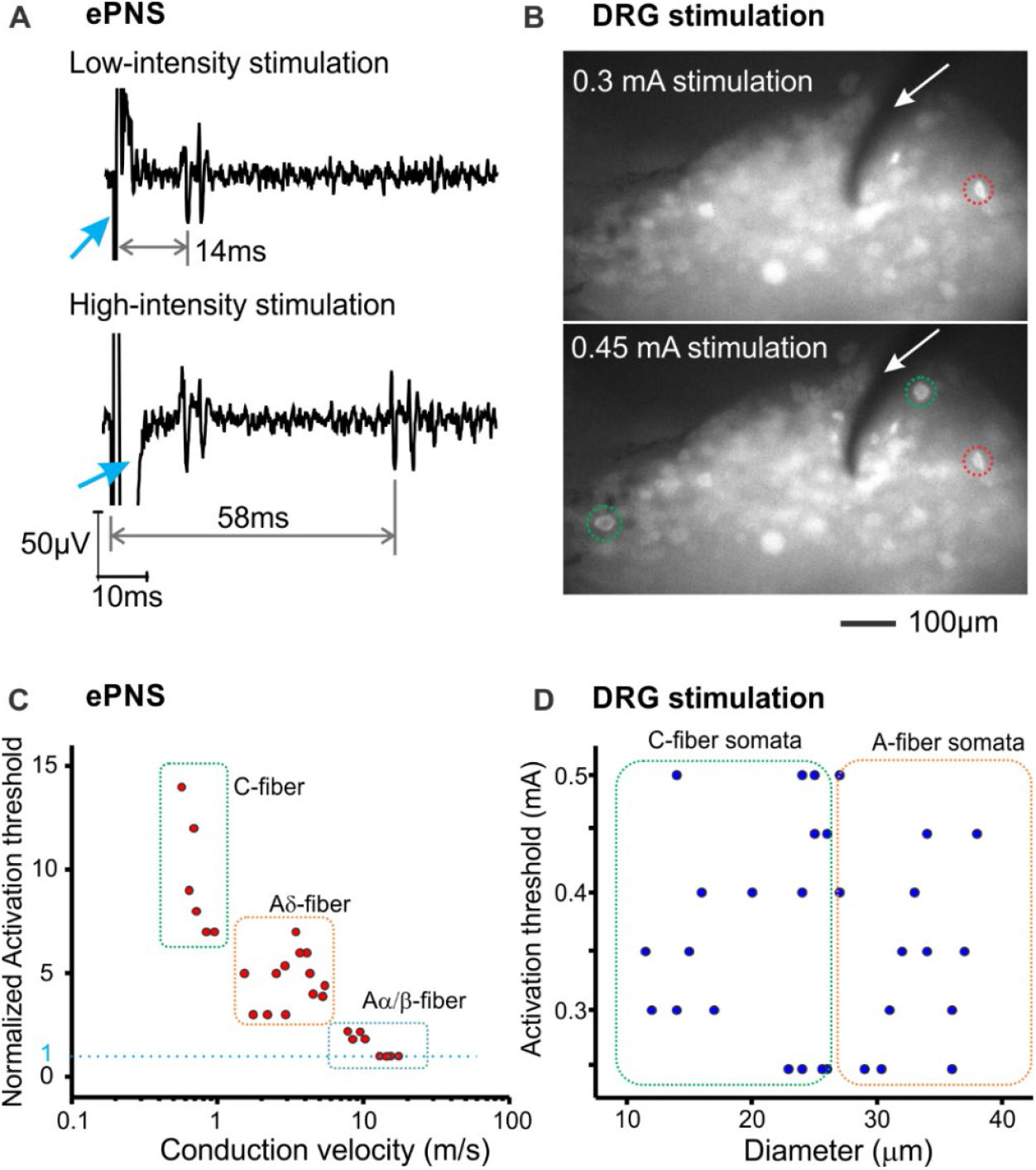
Differential activation of afferent populations by ePNS and DRG stimulation. (A) Representative single-fiber recordings from one split sciatic nerve filament showing lower stimulus threshold for activating Aδ-fibers (14 ms conduction delay) than for activating C-fibers (58 ms conduction delay). Arrow: stimulus artifact. (B) Representative GCaMP6f image recordings demonstrating activation thresholds of neurons with different diameters. White arrow: stimulating electrode; Red dashed circle: a small-diameter neuron (Φ17 μm) with 0.3 mA threshold, green dashed circles: two large-diameter neurons (Φ34 μm, Φ28 μm) with 0.45 mA threshold. (C) Normalized activation thresholds of peripheral axons with different conduction velocities by ePNS, measured by single-fiber recording. (D) Activation thresholds of afferents with different soma diameters by DRG stimulation, measured by GCaMP6f imaging.

### Differential distribution of NaV1.6 in A- and C-fiber afferents

By analyzing published scRNA-seq datasets ^18, 19^, we found that NaV1.6 mRNA is widely expressed in both A- and C-type afferent neurons (**Fig. 2**). We then performed immunohistological staining for NaV1.6 on fluorescently labeled afferent tissues from VGLUT2-Cre mice, labeled either via intrathecal delivery of AAV9-floxed-ChR-EYFP or by the tdTomato reporter line, to assess the distribution of NaV1.6 protein in afferent somata within the DRG, peripheral axons in the sciatic nerve, and nerve endings embedded in the bladder wall. Co-staining of NaV1.6 (red fluorescence) and afferent structure (green fluorescence) in DRG sections indicates clustering of NaV1.6 in the stem axon near the somata (**Fig. 4A**). Neurons were classified as large- and small-diameter classes following the same criteria as in the GCaMP6 study in **Fig. 3**. NaV1.6 clustering in the stem axon was observed in both large- and small-diameter neurons (**Fig. 4A**). Distance from the soma to the first NaV1.6 cluster along the stem axon was measured and compared between the large- and small-diameter groups and reported in **Fig. 4B**. The average distance measured in large-diameter neurons (n = 6, 7.51 ± 1.55 μm) was slightly greater than at in the small-diameter counterparts (n = 36, 6.42 ± 0.75 μm), but the difference was not statistically significant (t-test, p = 0.58).

**Figure 4.**
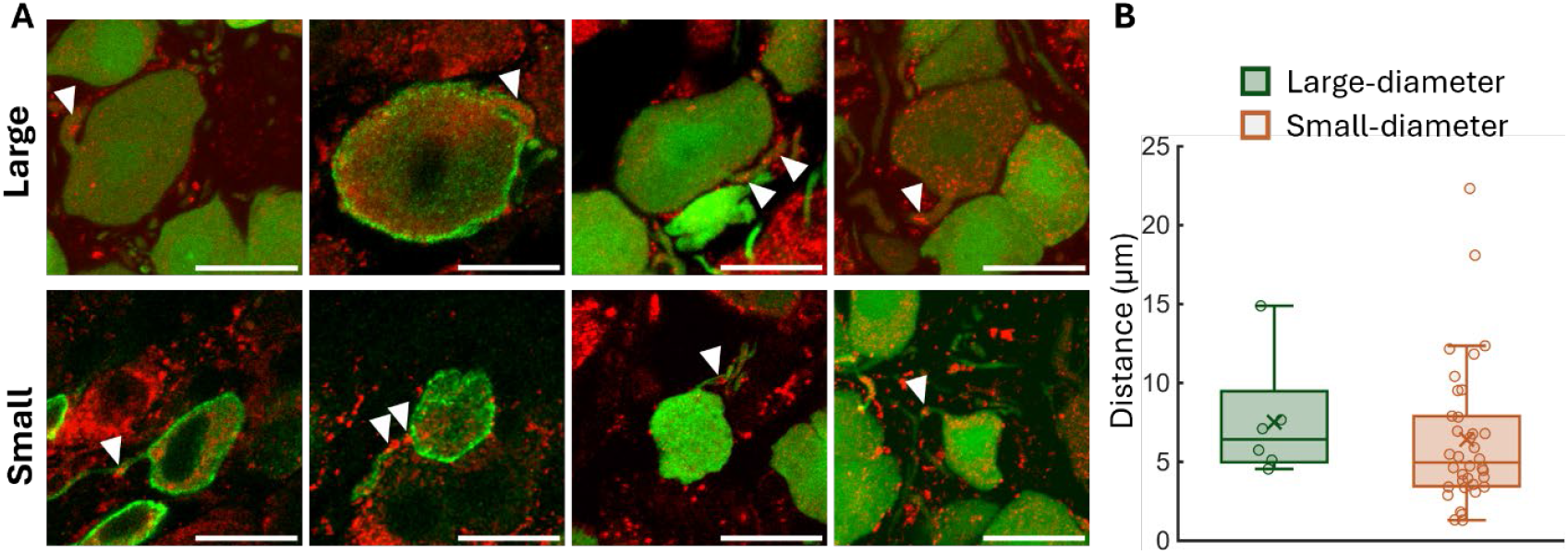
Both large- and small-diameter DRG neurons show clustering of NaV1.6 in stem axons near the somata. (A) Representative DRG sections co-stained for NaV1.6 (red) and afferents (green) showing NaV1.6 clusters (indicated by white arrow heads) in the stem axon of both large- and small-diameter neurons. Scale bar: 20 μm. (B) Distance between the somata and the first NaV1.6 clustering in the stem axon. There was no significant difference between the large- and small-diameter groups (t-test, p = 0.58).

Sciatic nerve with sparsely labeled afferent axons by intrathecal AAV9 injection were co-stained with NaV1.6 antibody as shown in **Fig. 5**. The clustering of NaV1.6 staining was detected in both Aβ-fiber (Φ > 4.5 μm) and Aδ-fiber axons (1.5 μm < Φ < 4.5 μm), whereas no clustering was observed in C-fiber axons (Φ < 1.5 μm). Clustering density was quantified as the number of distinct NaV1.6 clusters per 100 μm of traced sciatic axons. As shown in **Fig. 5B**, NaV1.6 clustering density was significantly higher in Aβ-fiber axons (n = 4, 5.60 ± 0.65 clusters/100 μm) than in Aδ-fiber axons (n = 9, 1.30 ± 0.16 clusters/100 μm; t test, p = 0.003).

**Figure 5.**
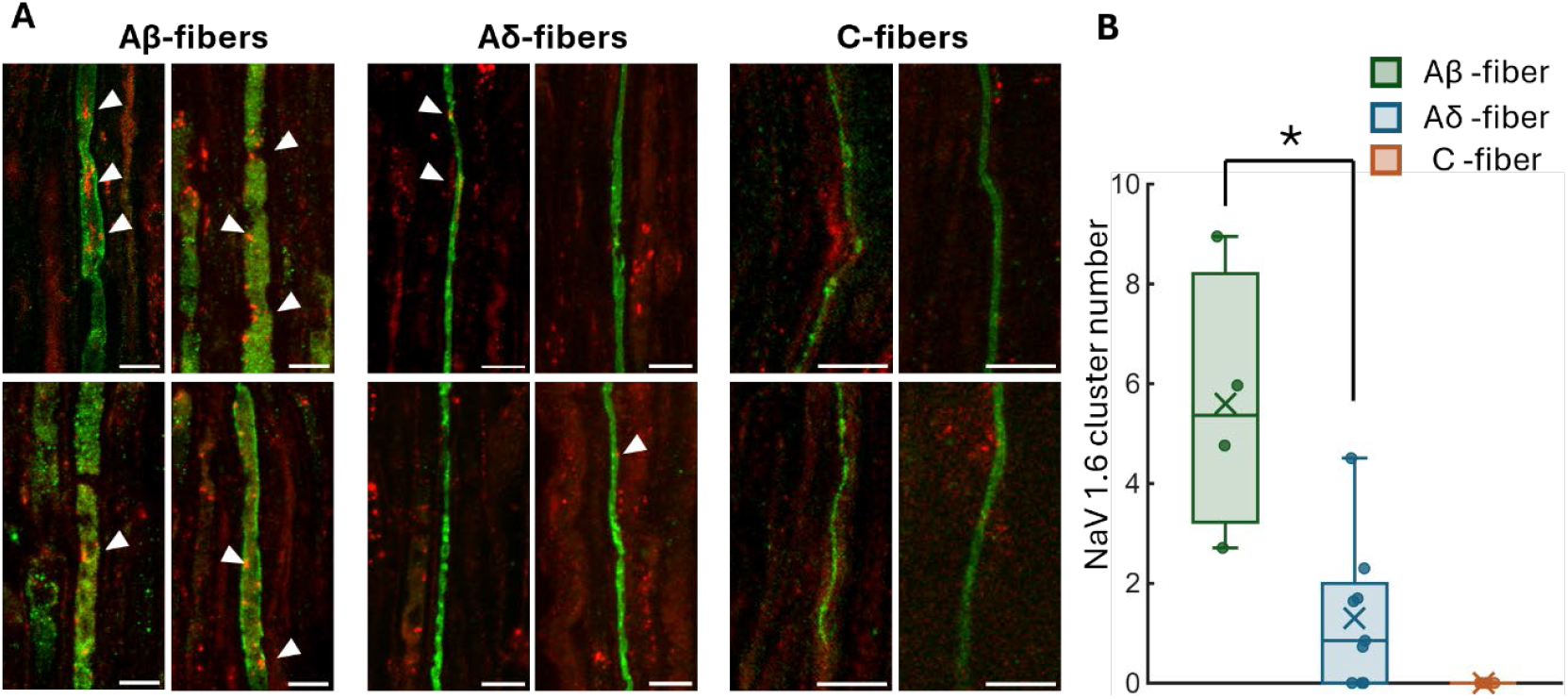
NaV1.6 clustering is present in A-fiber axons but absent in C-fiber axons of the sciatic nerve. (A) Representative immunostaining of NaV1.6 in sciatic nerve sections with sparsely labeled afferents following intrathecal AAV9 injection. Arrowheads indicate the presence of NaV1.6 clusters in Aβ- (Φ > 4.5 μm) and Aδ-fiber axons (1.5 μm < Φ < 4.5 μm), but not in C-fiber axons (Φ < 1.5 μm). (B) NaV1.6 cluster density (clusters per 100 μm) is significantly higher in Aβ-fiber axons than in Aδ-fiber axons (mean ± SEM; t test, p = 0.003). Scale bar: 10 μm.

Visceral afferents in the bladder, which consist predominantly of unmyelinated C-fiber afferents ^36^, were selected to assess NaV1.6 clustering in afferent nerve endings. Bladder tissue sections from VGLUT2-Cre/tdTomato mice, in which afferent axons and terminal endings were fluorescently labeled, were immunostained with a NaV1.6 antibody. Analysis of Z-stack images from 20 bladder sections (20 μm thickness) revealed colocalization of NaV1.6 with afferent axons in 5 images (17%), which consists of putative afferent terminal endings. Representative bladder afferent endings exhibiting NaV1.6 clustering are shown in **Figs. 6A-B**. In most bladder sections (83%) as shown in **Figs. 6C-D**, no NaV1.6 clustering was detected, likely due to the lack of terminal endings in the field of view.

**Figure 6.**
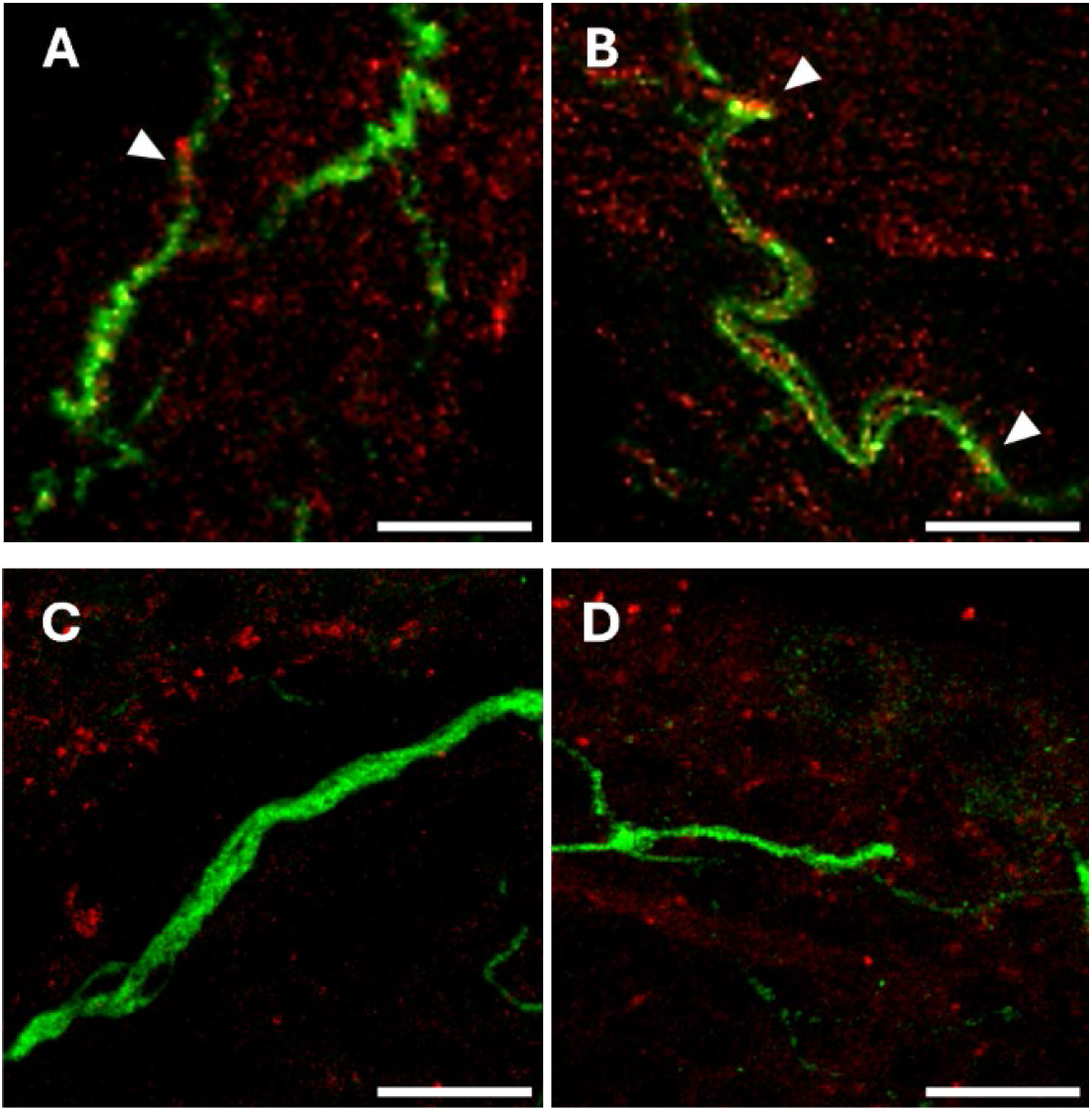
NaV1.6 clustering in afferent nerve endings in the bladder. (A-B) Immunostaining of bladder tissue sections from VGLUT2/tdTomato mice showed the presence of NaV1.6 clustering, in some bladder afferent axons (putative nerve terminals) as indicated by white arow heads. (C–D) Non-terminal axons of bladder afferents lack detectable NaV1.6 staining. Scale bar: 20 μm.

### Critical role of NaV1.6 clustering in lowering the activation threshold of C-fiber afferents through NEURON modeling

A C-fiber afferent was simulated in the NEURON simulation environment as illustrated in **Fig. 1C**, consisting of a soma and stem, peripheral and central axons. A 4-μm-long NaV1.6 clustering zone was positioned in the stem axon, 6.4 μm from the soma. Extracellular DRG stimulation evoked AP near the soma, which propagated toward both the peripheral and central axons (**Fig. 7A**). In contrast, extracellular ePNS initiated APs near the peripheral terminal, which then propagated to the soma and central axon (**Fig. 7B**). The stimulus threshold required to evoke APs during DRG stimulation depended strongly on NaV1.6 clustering in the stem axon. Specifically, the threshold decreased progressively with increasing maximum NaV1.6 conductance within the clustering zone, to a much greater extent than comparable increases in either NaV1.7 or NaV1.8 conductance (**Fig. 7C**). In comparison, the threshold for ePNS is 0.07 mA, similar to the threshold of DRG stimulation in the absence of NaV1.6 clustering (0.065 mA; **Fig. 7C**). Spatial voltage plots further demonstrated that AP initiation during DRG stimulation originated within the NaV1.6 clustering zone (**Fig. 7D**).

**Figure 7.**
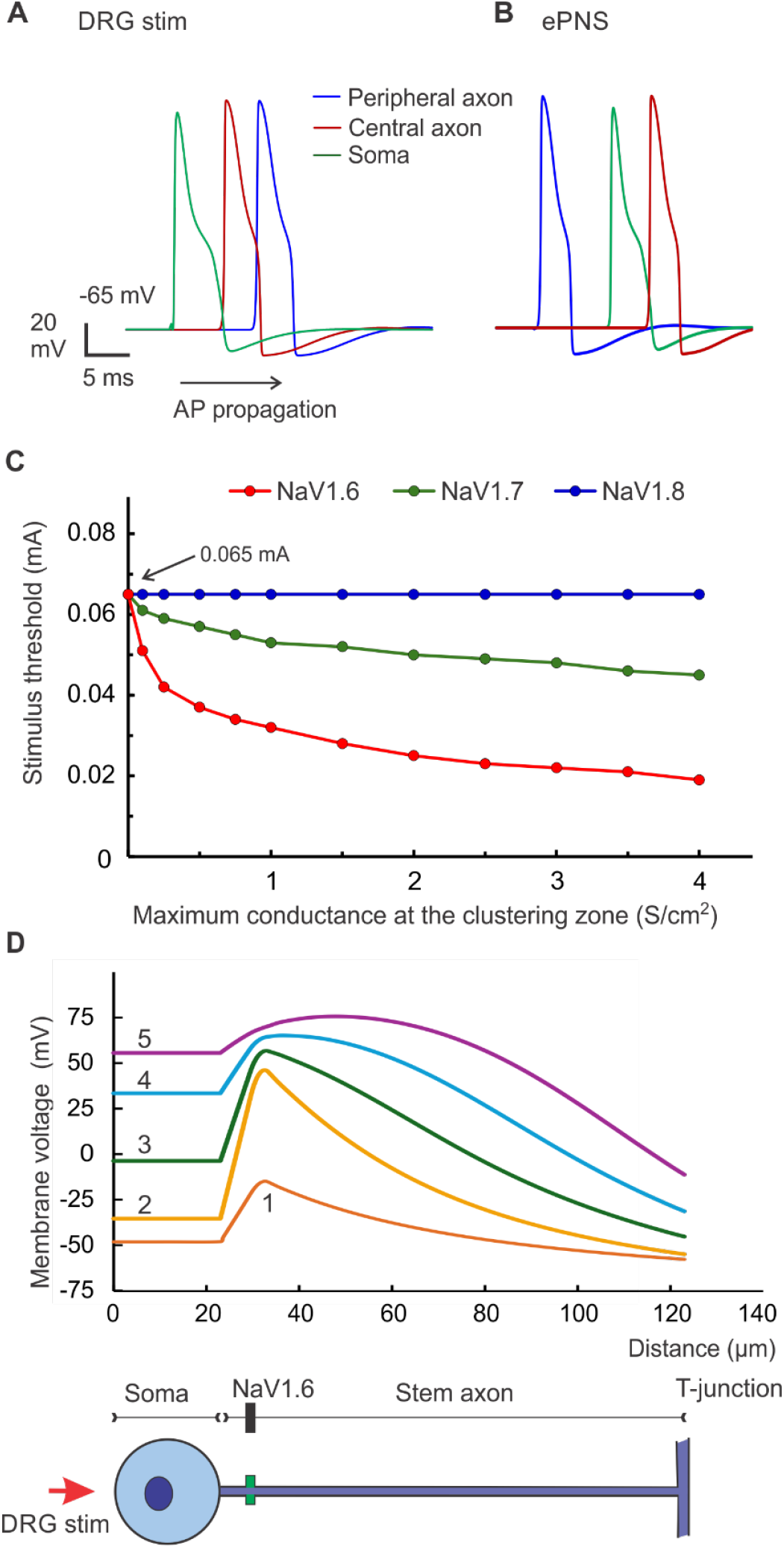
NEURON simulation of ePNS and DRG stimulation of a C-fiber afferent. (A) Action potentials evoked by DRG stimulation initiates close to the soma and propagates bi-directionally to both peripheral and central axons. (B) Action potentials evoked by ePNS of the peripheral axon propagates to soma and central axon. (C) The clustering of NaV1.6 in the stem axon strongly decreased the threshold of DRG stimulation to evoke APs. In contrast, comparable increases in NaV1.8 conductance within the clustering zone had no effect on the threshold, whereas increases in NaV1.7 conductance produced only a modest reduction. (D) Spatial plot along the soma and stem axon to reveal the site of spike initiation at the NaV1.6 clustering zone. Five spatial membrane voltage traces have 0.05 msec intervals (from 1 to 5).

## Discussion

This study systematically characterized the activation thresholds of A- and C-fiber axons by electrical peripheral nerve stimulation (ePNS) and dorsal root ganglion (DRG) stimulation using high-resolution single-fiber electrophysiological recordings and optical GCaMP6f recordings from individual DRG neurons. Single-fiber recordings from a split nerve filament often enable simultaneous recording from multiple axons with different conduction velocities; the large differences in conduction delays allow straightforward separation of spikes arising from distinct afferents ^25, 26^. From the same split nerve filament, stimulus thresholds required to evoke fast-conducting A-fibers and slow-conducting C-fibers were measured and directly compared under identical electrical stimulation conditions. We found that the average threshold required to activate C-fiber axons by ePNS was approximately 10-fold higher than that required to activate fast-conducting Aα/β-fiber axons, highlighting the challenge of selectively activating C-fiber axons using ePNS. These results from single-fiber recordings are consistent with prior whole-nerve compound action potential (CAP) recordings, in which substantially higher stimulus intensities are required to evoke slow-conducting C-fiber volleys than fast-conducting A-fiber volleys ^37^. It should be noted that the monopolar configuration used in our single-fiber recordings produces a sizable stimulus artifact that obscures evoked spikes with extremely short conduction delays (<2 ms). Consequently, Aα/β-fiber axons with conduction velocities greater than 12 m/s were not analyzed; these fibers likely have even lower activation thresholds. Therefore, the observed ∼10-fold difference in activation thresholds between C- and Aα/β-fiber axons by ePNS is likely an underestimate.

To systematically assess afferent activation thresholds by DRG stimulation, we implemented GCaMP6f recordings from intact DRG, an optical approach we previously established for evaluating afferent excitability ^27, 38^. GCaMP6f exhibits rapid kinetics in response to intracellular calcium elevation and has been validated for detecting individual action potentials in DRG neurons ^27, 38^ as well as in trigeminal ganglion neurons ^39^. Although GCaMP6f recordings do not directly determine axonal conduction velocity, neuronal soma size provides a reliable proxy for myelination status. Patch-clamp electrophysiology and immunohistological studies have established a strong correlation between DRG neuron size and myelination; mouse DRG neurons with diameters <27 μm are generally considered unmyelinated C-fiber afferents ^33, 34^. In contrast to the large difference in activation thresholds observed with ePNS, we found that DRG stimulation over a narrow amplitude range (0.25–0.5 mA) evoked activity in both small- and large-diameter DRG neurons, corresponding to putative C- and A-fiber afferents. This contrasts sharply with our current report of over 10-fold difference in activation thresholds between C- and A-fiber axons during ePNS. Among neurons activated by DRG stimulation, 57% were classified as C-fiber afferents, a proportion slightly lower than the anatomically estimated C-fiber population in rodent DRG (65 – 94%) ^40, 41^. To minimize photobleaching, experimental duration was limited by adjusting stimulus amplitude in 0.05 mA increments. To avoid confounding effects of suprathreshold stimulation, which can recruit excessive numbers of neurons, the maximum stimulus amplitude was capped at 0.5 mA, and analysis was restricted to neurons located near the stimulation electrode. It is possible that higher stimulus intensities could recruit additional myelinated A-fiber afferents located farther from the electrode, further reducing the apparent proportion of activated C-fiber neurons. Nonetheless, these data strongly indicate that a subset of C-fiber afferents exhibits activation thresholds comparable to those of A-fiber afferents during DRG stimulation, supporting the conclusion that DRG stimulation activates both myelinated and unmyelinated afferents, in contrast to the predominantly myelinated axon activation observed with ePNS.

The presence of NaV1.6, a sodium channel subtype characterized by low activation thresholds and persistent and resurgent currents ^20^, is typically associated with neuronal hyperexcitability and rapid repetitive firing, which are features seemingly inconsistent with C-fiber nociceptors exhibiting high response thresholds and low firing rates. Paradoxically, our analysis of published scRNA-seq datasets revealed high NaV1.6 expression in DRG neurons expressing molecular markers of unmyelinated C-fiber afferents ^19^. Within the C-fiber population, NaV1.6 expression is high in peptidergic nociceptors expressing CGRP and TRPV1 and low in non-peptidergic nociceptors expressing P2X3. This transcriptional profile was corroborated by patch-clamp electrophysiological studies demonstrating that NaV1.6 contributes approximately 34% of the tetrodotoxin-sensitive sodium current in small-diameter DRG neurons ^42^. Enrichment of NaV1.6 in TRPV1-positive DRG neurons is further supported by elevated NaV1.6 mRNA levels in colonic DRG neurons ^18^, which predominantly comprise slow-conducting C-fiber afferents ^24^. Also, TRPV1 is highly expressed in visceral afferents ^43, 44^ and plays a critical role in visceral nociception ^45, 46^. Despite these observations, the physiological and pathophysiological role of NaV1.6 in C-fiber nociceptors remains incompletely understood. Conditional deletion of NaV1.6 in NaV1.8-positive small-diameter DRG neurons does not significantly alter evoked pain behaviors in mouse models of chronic neuropathic or inflammatory pain ^42^. In the present study, immunohistological analyses revealed an absence of NaV1.6 in putative C-fiber axons with small diameters in the sciatic nerve, consistent with prior reports ^42, 47^. In contrast, we observed prominent clustering of NaV1.6 in bladder afferent endings, consistent with our previous report of NaV1.6 clustering in colonic afferent endings ^23^. Combined pharmacological experiments and NEURON modeling indicate that NaV1.6 plays a critical role in tonic firing at the spike initiation zone of C-fiber afferent endings ^23^. To our knowledge, this study provides the first evidence of NaV1.6 clustering within the stem axons of small-diameter DRG neurons, representing putative C-fiber afferents. Similar clustering was also observed in stem axons of large-diameter DRG neurons. We previously reported the rheobase for DRG stimulation and proposed that spike initiation during DRG stimulation occurs at the stem axon rather than the soma ^16^, a conclusion further supported by the present identification of NaV1.6 clustering in stem axons. Consistent with this interpretation, our NEURON simulations indicate that extracellular DRG stimulation initiates action potential at the NaV1.6 clustering zone within the stem axon.

Our NEURON model of a C-fiber afferent incorporated four sodium channel subtypes, NaV1.6, NaV1.7, NaV1.8, and NaV1.9, the combined expression of which is known to govern neuronal excitability based on prior patch-clamp studies. NaV1.1 was excluded because it is predominantly expressed in myelinated A-fiber afferents; its contribution to differential activation thresholds between A- and C-fibers during ePNS remains to be examined in future studies. We employed previously published modeling frameworks, including Markov-type state models that capture subtle yet functionally important differences in gating between NaV1.6 and NaV1.7 ^23^. NaV1.9 generates a persistent subthreshold sodium current and is expressed in both A- and C-fiber afferents, making it unlikely to account for their differential activation thresholds during ePNS. NaV1.7 and NaV1.8 are highly expressed in unmyelinated C-fiber axons ^48, 49^. However, they seem to have limited contribution to axonal excitability given that the activation thresholds of C-fiber axons are more than 10-fold higher than of A-fiber axons despite the much lower expression level of NaV1.7 and NaV1.8 in A-fiber axons. Our sensitivity analyses revealed that small changes in NaV1.6 conductance within the clustering zone of the stem axon substantially altered activation thresholds during DRG stimulation, whereas ePNS thresholds remained high in axons lacking NaV1.6 conductance.

## Conclusions

This study provides direct functional evidence for distinct activation thresholds of unmyelinated C-fiber afferents during DRG stimulation versus ePNS. Immunohistological analyses revealed differential expression of NaV1.6 in C-fiber afferent axons, with prominent clustering in stem axons within the DRG but absence in sciatic nerve axons, supporting more efficient activation of C-fiber afferents by DRG stimulation than by ePNS. Whether NaV1.6 clustering in stem axons is required for the reduced activation threshold of C-fiber afferents during DRG stimulation remains beyond the scope of the present study and will require future genetic knockdown approaches. Together, these experimental and computational findings strongly support a beneficial role for DRG stimulation in engaging additional afferent-blocking mechanisms to suppress pain, particularly through targeted modulation of TRPV1-positive C-fiber nociceptors via activity-dependent conduction block. Indeed, low-frequency DRG stimulation effectively blocks C-fiber–mediated visceral nociception ^16^. These findings suggest that DRG stimulation may provide superior therapeutic efficacy for chronic pain conditions driven by C-fiber sensitization, which clinically manifest as burning, spontaneous, and ongoing pain that constitutes the predominant complaints of many chronic pain patients.

## Acknowledgments

This work was supported by NSF CAREER Award 1844762 and NIBIB EB037604 grant to Dr. Feng.

